# Outer membrane vesicles from *Bacteroides fragilis* contain coding and non-coding small RNA species that modulate inflammatory signaling in intestinal epithelial cells

**DOI:** 10.1101/2025.06.25.661399

**Authors:** Aadil Sheikh, Colin Scano, Julia Xu, Tolulope Ojo, Jessica M. Conforti, Kayla L. Haberman, Bryan King, Alysia S. Martinez, Michelle Pujol, Juli Watkins, James Lotter, Emily L. Lin, Bernd Zechmann, Amanda Sevcik, Christie Sayes, Elyssia S. Gallagher, Steven P. Lang, Joshua Mell, Garth D. Ehrlich, Joseph H. Taube, K. Leigh Greathouse

**Affiliations:** Department of Biology, Baylor University, Waco, USA; Department of Chemistry & Biochemistry, Baylor University, Waco, USA; Center for Microscopy and Imaging, Baylor University, Waco, USA; Department of Environmental Sciences, Baylor University, Waco, USA; Department of Microbiology and Immunology, Drexel University, Philadelphia, USA; Department of Human Sciences and Design, Baylor University, Waco, USA

**Keywords:** Bacteroides fragilis, outer membrane vesicles, small RNA

## Abstract

Alterations to the community structure and function of the microbiome are associated with changes to host physiology, including immune responses. However, the contribution of microbe-derived RNAs carried by outer membrane vesicles (OMVs) to host immune responses remain unclear. This study investigated the role of OMVs and OMV-associated small RNA (sRNA) species from pathogenic and commensal *Bacteroides fragilis* (ETBF and NTBF respectively) in eliciting different immune responses from intestinal epithelial cells. To distinguish the differences in the sRNA profiles of the two strains and their OMVs, RNA-seq, qRT-PCR, and northern blotting were conducted to identify enrichment of discrete sRNA species in OMVs, which were also differentially expressed between the two strains. Specifically, both coding and non-coding RNAs were enriched in OMVs from NTBF and ETBF, with BF9343_RS22680 and BF9343_RS17870 being significantly enriched in ETBF OMVs compared to NTBF. To understand the effects of OMVs on pattern recognition receptors, reporter cells of Toll-like receptor (TLR) activation were treated with OMVs, demonstrating activation of TLRs 2, 3, and 7. Treatment of Caco-2 and HT29-MTX cells with OMVs demonstrated increased expression of IL-8. Surprisingly, we discovered that degradation of RNase-accessible RNAs within ETBF OMVs, but not NTBF OMVs, resulted in vesicles with enhanced capacity to stimulate IL-8 expression, indicating that these extravesicular RNAs exert an immunosuppressive effect. This suggests a dual role for OMV-associated RNAs in modulating host immune responses, with implications for both bacterial pathogenesis and therapeutic applications.

**Graphical Abstract**

## Introduction

Colorectal cancer (CRC) is the third most commonly diagnosed cancer in the United States and represents the second most common cause of global cancer-related mortality.^1,2^ Chronic intestinal inflammation is a key factor driving CRC pathogenesis,^3,4^ as exemplified by patients with inflammatory bowel disease (IBD) who face an elevated risk of developing colitis-associated CRC (CAC).^5–7^ Over time, persistent inflammation associated with IBD disrupts the intestinal barrier, promotes epithelial injury, and fosters an environment conducive to tumorigenesis..^8–11^

A crucial determinant of this inflammatory milieu is the composition and functional state of the gut microbiome. The intricate community of bacteria, viruses, and fungi within the gut typically exists in a symbiotic relationship with the host. However, alterations in the microbial community structure can lead to sustained inflammation ^12–14^, predisposing to CRC development.^15,16^ Among the microbial constituents implicated in CRC pathogenesis is *Bacteroides fragilis*, which is commonly found in the gut as a non-enterotoxic (NTBF) strain that generally supports immunotolerance.^17,18^ In contrast, enterotoxigenic *B. fragilis* (ETBF) produces a toxin (BFT or fragilysin) that cleaves E-cadherin in tight junctions, thereby compromising the epithelial barrier and promoting pro-inflammatory signaling.^19,20^ Although BFT is well-characterized, other bacterial factors also appear to drive inflammation ^21,22^, and their precise modes of secretion and delivery to host cells remain under investigation.

One avenue for bacterial-host communication is the release of outer membrane vesicles (OMVs).^23,24^ These 50–250 nm vesicles, shed by Gram-negative bacteria including *B. fragilis*, carry a variety of biologically active components—proteins, lipids, lipopolysaccharides (LPS), peptidoglycans (PGs), and nucleic acids ^25–27^—that can directly interact with host cells.^28–30^ OMVs have been shown to activate pattern recognition receptors (PRRs), such as the Toll-like receptor (TLR) family, leading to downstream immune cascades. TLRs recognize pathogen-associated molecular patterns (PAMPs) and can be categorized as extracellular and intracellular (endosomal) entities.^28^ Extracellular TLRs recognize lipooligosaccharides (LOS), LPS, PGs, and flagellin, among other ligands, whereas endosomal TLRs bind to foreign DNA, dsRNA, and ssRNA taken up by host cells.^28,31–34^ TLR activation typically results in a cascade that leads to cytoplasmic phosphorylation of the NFκB and IRF transcription factors, which can translocate to the nucleus and induce the expression of pro-inflammatory (e.g., TNF-α, IL-1*β*, IL-6, and IL-8) or anti-inflammatory (e.g., IL-10) cytokines. While most research on OMV-mediated TLR activation has focused on canonical bacterial ligands such as LPS and PGs, recent studies highlight OMV-associated RNAs as potential drivers of host immune responses.^25,35–41^

Despite growing awareness of the importance of vesicular RNA, major questions remain regarding the specific RNA sequences that become enriched in OMVs, and how these RNAs may influence host cells differently when derived from pathogenic versus commensal strains. To address these gaps, we profiled OMVs from NTBF and ETBF, assessing their size, protein enrichment, and RNA cargo. We identified distinct RNA species—some protected from, and others susceptible to, RNase degradation—that appear to modulate the inflammatory response. Notably, RNase-sensitive OMV-associated RNAs conferred an immunosuppressive effect, reducing the ability of OMVs to induce IL-8 expression. By revealing an unexpected role for vesicular RNAs in shaping immune responses, our study provides insight into how bacterial RNA cargo can either exacerbate or temper inflammation in the intestinal environment. This knowledge holds promise for guiding the development of OMV-based therapeutics aimed at modulating gut inflammation and mitigating CRC risk.

## Materials and Methods

### Bacteria strain and culture preparation

Enterotoxigenic *Bacteroides fragilis* (ETBF; 86-5443-2-2) and non-toxigenic *Bacteroides fragilis* (NTBF; NCTC 9343) were provided as a gift by Dr. Cindy Sears at Johns Hopkins University. Bacteria were cultured in brain heart infusion (BHI) broth under anaerobic conditions overnight at 37°C until an OD_600_ reading of 0.8 – 1.0 was achieved, which is characterized as exponential phase for this bacterium. All OMV isolations proceeded using the same OD_600_ for the exponential phase.

### Ultracentrifugation isolation of outer membrane vesicles

One liter of NTBF and ETBF cultures were spun down at 8000 x g for 15 minutes at 4°C and the resulting supernatant was filtered through 0.45 μm vacuum filter (Thermo Scientific, 168-0045). The filtered supernatant was ultracentrifuged at 100, 000 x g for 2 hours at 4°C (Beckman Coulter 70 Ti), and pelleted vesicles were resuspended in PBS. All preparations were either immediately used for downstream experiments or kept no more than 24 hrs at 4°. Vesicular sample concentrations were determined with the Pierce BCA Protein Assay Kit (Thermo Scientific, 23227).

### Column isolation and concentration of outer membrane vesicles

OMVs were isolated from NTBF and ETBF using the ExoBacteria OMV Isolation Kit (System Biosciences, EXOBAC100A-1), which is a precipitation-free, ion-exchange chromatography system, as per the manufacturer’s instructions. Briefly 250 mL of ETBF and NTBF cultures were spun down at 8000 x g for 15 minutes at 4°C and the resulting supernatant was filtered through 0.45 μm vacuum filter (Thermo Scientific, 168-0045). Filtered supernatant was run through the ExoBacteria OMV isolation column, and each column yielded 1.5 mL of eluted OMVs. 5 columns were used per culture resulting in 15 mL of each strain. The resulting solution of OMVs was concentrated using centrifugal concentration tubes (Thermo Scientific, 88532) at 5500 x g for 15 minutes. Vesicular sample protein concentrations were determined by disruption using RIPA buffer and quantification with the Pierce BCA Protein Assay Kit (Thermo Scientific, 23227).

### OMV protein characterization

OMV samples were diluted in SDS buffer to concentrations of 1500 μg/mL, 1000 μg/mL, 500 μg/mL and 200 μg/mL. All samples were run on a 10% SDS-PAGE gel (Bio-Rad, 4568033) at 175V. The gel was subjected to silver staining using the Pierce Silver Stain Kit (Thermo Scientific, 24612) as per the manufacturer’s instructions.

### OMV size profiling

Nanoparticle tracking analysis was conducted using the Exoid (Izon Science Limited) with default parameters.

### Transmission Electron Microscopy

Concentrated OMVs were prepared for TEM imaging by staining with uranyl acetate and osmium tetroxide separately, as described previously ^42^, then co-staining with osmium tetroxide and uranyl acetate. For uranyl acetate, copper grids were incubated on 10 *μ*L drops of OMVs for 5 minutes, followed by two washes on DI water drops for 2.5 minutes each and finally stained with 2% uranyl acetate for 1 minute. For the osmium tetroxide and uranyl acetate co-stain, 10 *μ*L of OMVs mixed with an equivalent volume of 4% osmium tetroxide for 10 minutes on copper grids followed by staining with 2% uranyl acetate for 1 minute. All grids had excess liquid blotted off with filter paper and were allowed to dry overnight. The grids were then imaged with a TEM (JEM-1010, JEOL Inc., Tokyo, Japan), a minimum of 7 images per test condition were obtained.

### Mass Spectrometry

Proteins were identified using liquid chromatography with tandem mass spectrometry (LC-MS/MS). Proteins from NTBF and ETBF OMVs were excised from the 25 kDa, 40 kDa, 55 kDa, 70 kDa, and 100 kDa bands of SDS-PAGE gels and prepared by in-gel digestion. LC-MS/MS data was searched against a Bacteroides fragilis reference proteome to identify proteins. A full description of the proteomics methodology can be found in the **Supplemental Information**.

### Small RNA isolation

Matched samples of RNA isolated from whole bacteria and corresponding OMVs were isolated in three individual preparations. RNA samples were isolated using the GeneJET RNA Purification Kit (Thermo Scientific, K0732) as per the manufacturer’s instructions for bacteria RNA isolation. RNA quality was assessed with a Bioanalyzer (Agilent Technologies, Santa Clara, CA, USA) and RNA concentrations were determined using the Qubit3 RNA HS Assay Kit (Invitrogen, Q32852).

### RNase protection assay and total RNA isolation

Concentrated OMVs were incubated with 10 μg/mL of RNase A at 37°C for 1 hour. The solution was then treated with RNA*secure* RNA Inactivation Reagent (Invitrogen; AM7006) for 10 minutes. Three matched preparations of whole cell pellets, RNase-treated OMVs and untreated OMVs were subjected to hot phenol RNA extraction. Whole cell pellets were treated in 1.5 mL of 0.5 mg/mL of lysozyme in 10 mM of Tris-HCl solution and incubated at 37°C for 5 minutes. For each 1.5 mL of whole cell and OMV samples, 150 μL of 10 % SDS and 30 μL of 7.5 M ammonium acetate was added followed by equal amounts of phenol:chloroform:isoamyl alcohol (25:24:1; pH ∼5.2) which was then vortexed until the solution was emulsified. Samples were incubated at 65°C for 15 minutes with a brief vortex every 5 minutes and then chilled on ice for 5 minutes. Samples were centrifuged at 15, 000 x g for 10 minutes at 4°C, and the aqueous layer was collected before an equal amount of chloroform was added. Samples were centrifuged again at 15, 000 x g for minutes at 4°C and the aqueous layer was removed. 100 μL of 7.5 M ammonium acetate and 2X chilled 95% ethanol were added to the mixture, vortexed, and centrifuged at 15, 000 x g for 20 minutes at room temperature. The supernatant was decanted, and 80% ethanol was added to wash the RNA pellet prior to centrifugation at 15, 000 x g for 20 minutes for 4°C. Lastly, the supernatant was decanted and allowed to air dry briefly before resuspension in 100 μL of ultrapure water.

### RNA sequencing and analysis of *Bacteroides fragilis* small RNAs

Small RNA extractions were quantified by Quant-iT RiboGreen RNA assay (Life Technologies, Thermo-Fisher) and assessed for purity using a Thermo Scientific Nanodrop 2000 spectrophotometer. To create indexed sequencing libraries, the Takara Biosciences (formerly Clontech) SMARTer smRNA-Seq kit for Illumina (catalog 635029) was used, which is suitable for sequencing small RNAs from low-yield samples. Illumina adapter and index sequences were incorporated without ligation to ensure diverse small RNAs are represented. 50 ng of starting material was used for each library.

Resulting libraries were quantified using Biotium Accuclear 7 DNA standards High Sensitivity assay (fluorometric quantitation) and the size distribution determined by Agilent Bioanalyzer using the High Sensitivity DNA assay. Fragment molarity and samples were normalized to 1.0 nM before pooling for sequencing at loading concentration of 1.8 pM on the Illumina NextSeq 500 using a High output 75 cycle reagent kit v2. Sequence reads were then demultiplexed using bcl2fastq2 before subsequent bioinformatic analysis.

Reads from all samples were trimmed and aligned to each *B. fragilis* genome (ETBF: 86-5443-2-2, NTBF: NCTC 9343) using the STAR aligner with default parameters. RNA enrichment analysis and visualization was completed using R (version 4.2.1 – “Funny-Looking Kid’) and RStudio (version 2022.07.2 – “Spotted Wakerobin”). Data analysis and visualization was completed using tidyverse (version 1.3.2), DESeq2 package (version 1.4.2), ggplot2 (version 3.3.6), EnhancedVolcano (version 1.14.0), and pheatmap (version 1.0.12). Data tables provided as supplemental files 2 (ETBF aligned) and 3 (NTBF aligned).

### Read Coverage Analysis

RNA coverage from sequencing data was visualized with *Integrative Genome Viewer* (Version 2.8.12). Northern blot probes were created by taking the complementary sequence of the densest read regions of the identified genes and intergenic regions of interest.

### qRT-PCR of Bacterial RNAs

Total RNA (1 μg) was reverse-transcribed using the High-Capacity cDNA Reverse Transcription Kit (Applied Biosystems, 4368814) as per the manufacturer’s instructions. The qPCR was performed with diluted cDNA using primer sets for genes identified through RNA-seq analysis. PCR amplification of RNA was performed using *Power* SYBR Green PCR Master Mix (Applied Biosystems, 4367659). All qPCR experiments were run in technical quadruplicates and biological triplicates and a mean value was used for the determination of RNA levels. Images were generated using GraphPad Prism 9.

### Northern blotting of bacterial RNAs

Northern blots of total RNA samples were conducted as follows. Northern blot probes were created by taking the complementary sequence to regions-of-interest with high coverage in OMV-enriched RNA-seq data. Probes were labeled with DIG using the 2^nd^ generation DIG oligonucleotide 3’-end labeling kit (Roche, 03353575910) according to the manufacturer’s instructions and diluted to a concentration of 1 ng/mL in 10 mL of ULTRAhyb ultrasensitive hybridization buffer (Invitrogen, AM8670). 30 μg of RNA samples were loaded onto 15% Mini-PROTEAN Tris-borate EDTA (TBE)-Urea gel (Bio-Rad, 4566053) and run for 2 hours at 90 V in 1X TBE buffer. Transfer of RNA was completed in 1X TBE buffer onto a positively charged nylon membrane (Invitrogen, AM10102) for 3 hours at 60V in cold conditions. Blots were UV crosslinked and incubated in hybridization buffer for 1 hour before being probed overnight at 42°C. Probes were poured off and blots were washed twice in 2X SSC with 0.1% SDS buffer for 15 minutes, twice in 0.1X SSC with 0.1% SDS buffer with 5 minutes and 1X SSC buffer for 10 minutes. Blots were processed using the DIG wash and block buffer set (Roche, 11585762001) as per the manufacturer’s instructions and imaged using ChemiDoc MP Imaging System (Bio-Rad, 12003154) for 20 minutes.

### Cell lines

Caco-2 and HT29-MTX cells were received as a generous gift from Dr. Christie Sayes and HEK-Blue TLR cells were obtained from InvivoGen. All cell lines were cultured in Dulbecco’s Modified Eagle’s Medium (DMEM) (Corning Inc.; Kennebuck, ME, USA) supplemented with 10% fetal bovine serum (FBS) (Equitech-Bio Inc.; Kerrville, Texas, USA) and 1X antibiotics (Penicillin/Streptomycin, Lonza; Basel, Switzerland). HEK-Blue TLR cells were additionally supplemented with 100 μg/mL of Normocin (Invivogen, ant-nr-1), 1X HEK-Blue Selection (Invivogen, hb-sel), 100 μg/mL of Zeocin (Invivogen, ant-zn-05), and 30 μg/mL of blasticidin (Invivogen, ant-bl-05). Cell lines were tested for mycoplasma every two weeks. Incubation occurred at 37°C with 5% CO_2_.

### TLR reporter assays

HEK-Blue TLR reporter assays (Invivogen) were carried out as per the manufacturer’s specifications for specific TLRs. Cells were treated with 100 μg/mL and 10 μg/mL of ETBF and NTBF vesicles.

### RNA extraction and qRT-PCR

Caco-2 and HT29 cells were treated with 100 μg/mL and 10 μg/mL of unconcentrated NTBF and ETBF vesicles for 2 hours and 8 hours. Cells were lysed in the presence of Trizol Reagent. (Invitrogen, 15596026) and RNA was extracted following the manufacturer’s instructions. RNA (250 ng) was reverse-transcribed using the High-Capacity cDNA Reverse Transcription Kit (Applied Biosystems, 4368814) as per the manufacturer’s instructions. The qPCR was performed with diluted cDNA using primer sets for cytokine genes. PCR amplification of RNA was performed using *Power* SYBR Green PCR Master Mix (Applied Biosystems, 4367659). All qPCR experiments were run in technical quadruplicates and biological triplicates and a mean value was used for the determination of RNA levels. Images were generated using GraphPad Prism 9.

### Fluorescent Imaging of RNA and OMVs in Colonic Epithelial Cells

Staining of RNA and OMVs in colonic epithelial cells was performed as follows. Isolated OMVs from ETBF and NTBF strains were stained with SytoRNASelect (Thermo Fisher Scientific) to detect RNA. OMVs were incubated with SytoRNASelect at a final concentration of 5 μM in phosphate-buffered saline (PBS) for 30 minutes at 37°C in the dark. After staining, OMVs were washed twice by ultracentrifugation at 100,000 × g for 1 hour at 4°C to remove excess dye. For membrane labeling, RNA-stained OMVs were further incubated with Vybrant DiD (Thermo Fisher Scientific) at a final concentration of 1 μM for 30 minutes at 37°C in the dark. OMVs were washed again in PBS by centrifugation to remove unbound dye and resuspended in PBS for further use. Caco2 colonic epithelial cells were seeded on glass coverslips in 24-well plates at a density of 1 × 10⁵ cells per well and incubated in Dulbecco’s Modified Eagle Medium (DMEM; Thermo Fisher Scientific) supplemented with 10% fetal bovine serum (FBS), 1% non-essential amino acids, and 1% penicillin-streptomycin at 37°C in a humidified atmosphere with 5% CO₂. Cells were treated with stained OMVs (100 μg/mL) for 30 minutes at 37°C. Following treatment, cells were washed three times with PBS to remove unbound OMVs and fixed with 4% paraformaldehyde for 15 minutes at room temperature. Cells were then permeabilized with 0.1% Triton X-100 in PBS for 5 minutes and washed again with PBS. For nuclear staining, cells were incubated with DAPI (4’,6-diamidino-2-phenylindole, Thermo Fisher Scientific) at a concentration of 1 μg/mL in PBS for 5 minutes at room temperature in the dark. After staining, cells were washed three times with PBS. Coverslips were mounted onto glass slides using ProLong Gold Antifade Mountant (Thermo Fisher Scientific). For endoplasmic reticulum (ER) staining, cells were incubated with an anti-protein disulfide isomerase (PDI) antibody (1:200 dilution; Abcam) in PBS containing 1% bovine serum albumin (BSA) overnight at 4°C, followed by washing and incubation with Alexa Fluor 594-conjugated secondary antibody (1:500 dilution; Thermo Fisher Scientific) for 1 hour at room temperature in the dark. Stained cells were mounted and imaged using a Leica SP8 confocal laser scanning microscope, with image acquisition parameters optimized to avoid signal bleed-through between channels.

## Results

### Differentially enriched RNA contents between ETBF and NTBF OMVs and whole-cell

To test whether ETBF and NTBF package distinct RNA classes into their OMVs—our first prerequisite for a differential signalling mechanism—we isolated and sequenced small RNA from ETBF and NTBF OMVs, as well as whole-cell lysates of their reference bacterial cultures. Cross-alignment of reads to both ETBF and NTBF genomes was conducted to detect differentially enriched and unique sequences using DESeq2 analysis. We first characterized the proportion of gene coding sequence types in each of the samples (Fig. 1A/B). In the OMV samples, we found a significant enrichment of mRNAs and a depletion of rRNA and tRNA sequences, when compared to the whole cell (WC) samples, though both rRNA and tRNA reads were relatively abundant within OMVs.

**Figure 1:**
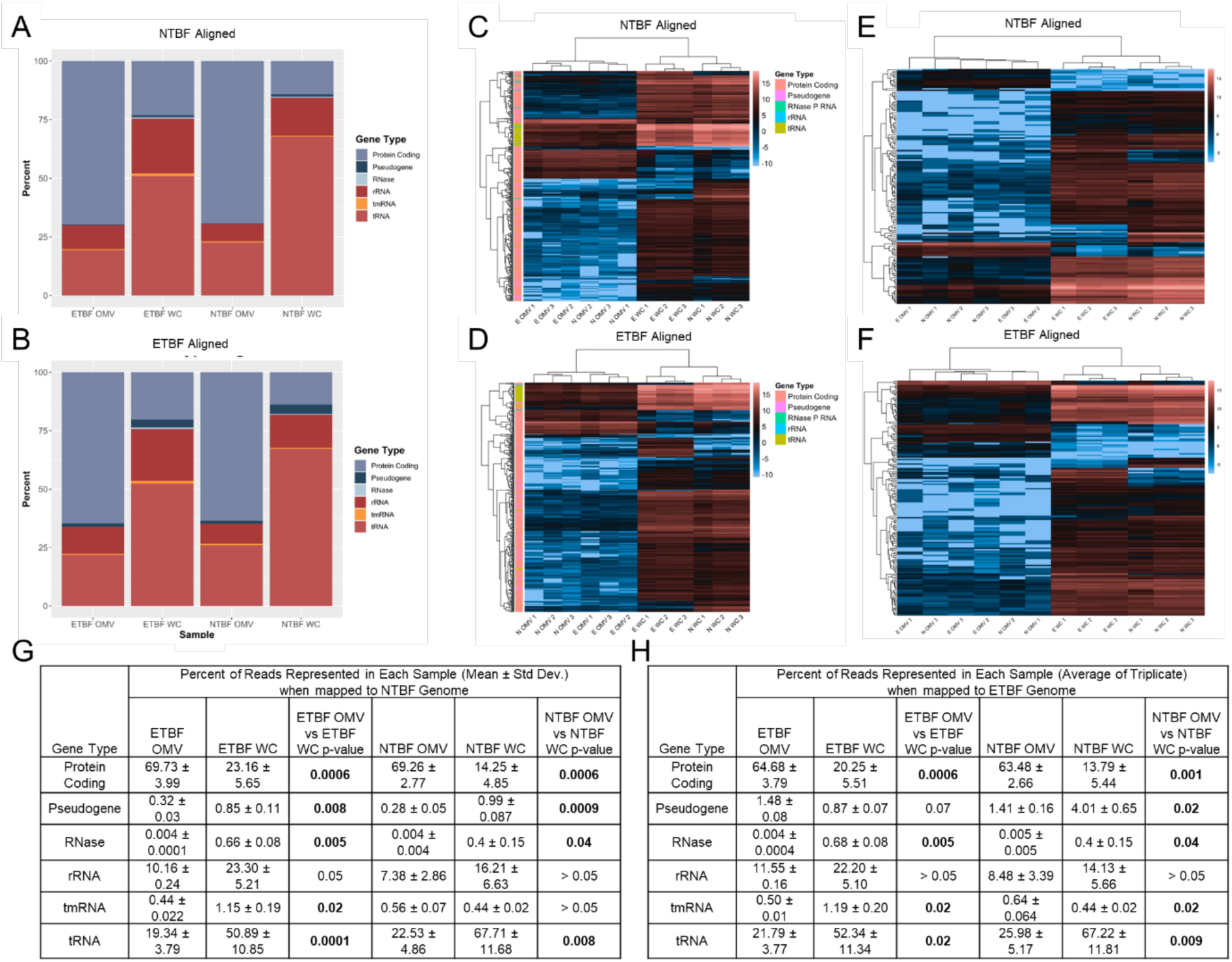
RNA sub-type enrichment is significantly different between ETBF and NTBF OMVs and whole cells. (A/B) the average proportion of sequencing reads mapped to the NTBF (A) and ETBF (B) genomes and represented by RNA class. Data are the mean of n = 3 biological replicates. (C/D) Unsupervised hierarchical clustering of DESeq2 analysis of gene coding regions using DESeq2 reveals enrichment of RNA sequences in OMVs and whole cell lysate (WC) samples when they are aligned to the NTBF (C) and ETBF (D) genomes. (E/F) Unsupervised hierarchical clustering of DESeq2 analysis of intergenic, non-coding regions aligned to the NTBF (E) and ETBF (F) genomes. E = ETBF and N = NTBF. (G/H) Proportion of transcript types represented in data aligned to NTBF genome (G) or ETBF genome (H). Statistical significance was determined using a paired t-test, p-value <0.05 significant.

Because selective loading of specific transcripts could underlie strain-specific host responses, we next profiled the top-enriched small RNAs in ETBF versus NTBF OMVs. We identified the most enriched RNA species in OMV samples by strain using DESeq2. For this, we first focused on sequences that mapped to known genes. RNAs loaded into OMVs by both strains of *B. fragilis* are more similar to each other than to the RNA retained in the whole cell, however sufficient differences remained to distinguish OMVs from each strain (Fig. 1C/D). While most sequences detected were depleted in OMVs relative to whole cell, a subset of protein-coding sequences were significantly enriched (Fig. 1C/D). As sRNAs may result from transcription of intergenic regions as well as from protein-coding or other functional categories, we next analyzed enrichment of reads that do not map to known genes. Similarly, analysis of intergenic reads confirms the loading of a subset of non-coding transcripts within both NTBF and ETBF OMVs (Fig. 1E/F).

Analysis of known transcripts present within OMVs successfully distinguished OMVs from intact cells, as well as distinguishing OMVs derived from NTBF and ETBF. To identify the specific RNAs responsible for these distinctions, we analyzed the RNA-seq data for statistical significance and fold change. While dozens of transcripts were depleted in OMVs (Fig. 2A-D), only two were significantly enriched in ETBF OMVs vs NTBF OMVs (Fig. 2E/F). One, BF9343_RS22680, partially encodes for a hypothetical protein and the other, BF9343_RS17870, partially encodes for a DUF4373 domain-containing protein.

**Figure 2:**
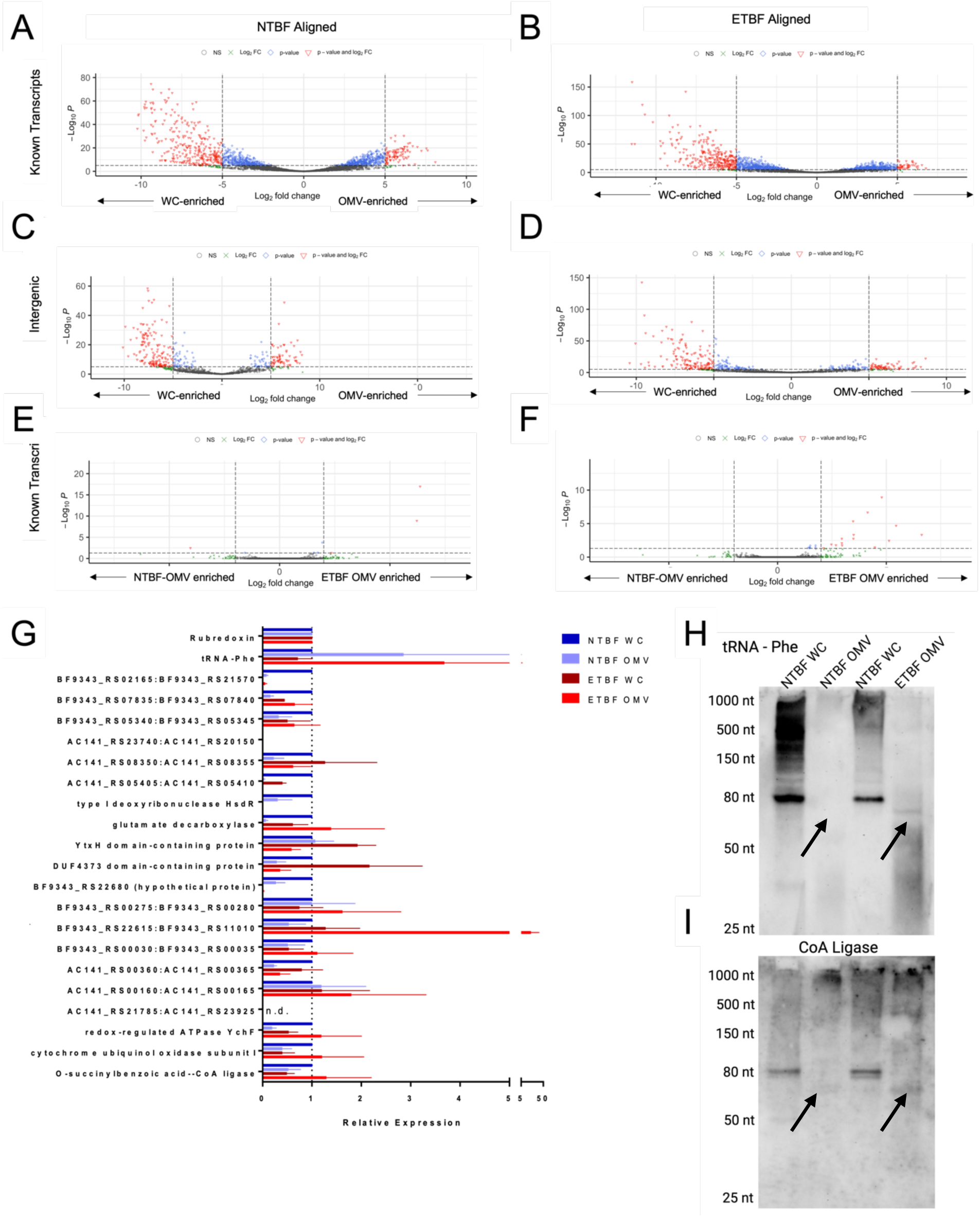
*B. fragilis* OMV RNA transcripts show differential abundance between ETBF and NTBF. (A/B) Enrichment of transcripts from established gene coding regions for combined OMV and whole cell lysate (WC) samples when they are aligned to the NTBF (A) and ETBF (B) genomes. (C/D) Enrichment of transcripts from intergenic regions for combined OMV and WC samples when they are aligned to the NTBF (C) and ETBF (D) genomes. (E/F) Enrichment of transcripts from established gene coding regions for OMV samples from ETBF and NTBF when they are aligned to the NTBF (E) and ETBF (F) genomes. All panels are derived from the same RNA-seq dataset; RNA transcript IDs are in the supplemental datasets. Limits of biological and statistical significance are log_2_FC was set at 5, and the adjusted p value was set to 10^-4^. (G) qRT-PCR for indicated known transcripts and intergenic regions. Normalized to rubredoxin and referenced to NTBF WC. n = 3 n.d. = not detected. (H/I) To confirm the size of tRNA (H) and CoA Ligase (I) transcripts, northern blot probes complimentary to the RNAseq reads were used and successfully detected transcripts in OMVs at lower length than in WC samples (denoted by arrows).

To validate both protein-coding and intergenic transcripts detected via RNA-seq, we selected a set of transcripts that were highly abundant or differentially enriched for validation with qRT-PCR and northern blotting. We targeted genes that met our enrichment criteria (Log_2_FC = 5, p-value = 10^-4^) and had high count numbers (∼2000 counts) from both established and intergenic transcripts. The two transcripts that were enriched in ETBF OMVs over NTBF OMVs (Fig. 2E) were also included. For normalization by qRT-PCR, we used the rubredoxin transcript, which was not differentially expressed between OMV and whole cell (WC) samples and was abundant in all samples. Twenty of 22 RNAseq-defined differentially expressed transcripts were verified by qRT-PCR (Fig. 2G). Notable enrichments include depletion of intergenic transcript AC141_RS05405:AC141_RS05410 in OMVs, NTBF-specific detection of type I deoxyribonuclease HsdR and intergenic transcript BF9343_RS22680, enrichment of glutamate decarboxylase in ETBF OMVs relative to NTBF OMVs, and enrichment of intergenic transcript BF9343_RS22615:BF9343_RS11010 in ETBF OMVs (Fig. 2G).

To validate the size of detected transcripts, we selected two RNAs for northern blotting. Probes for tRNA-Phe and for O-succinylbenzoic acid-CoA ligase (CoA ligase) were designed to be complementary to the highest coverage region of OMV-derived RNAseq read alignments (using the same RNA-seq data shown in Fig. 2A-F). The tRNA-Phe was detected as both a multi-tRNA transcript and a shorter, 80-nt fragment in NTBF and ETBF whole cell as well as in a ∼50-nt length version (arrow) in OMV samples (Fig. 2H). The CoA ligase transcript was detected at full length (>1000 nt) in all samples but also in a shorter 80-nt form in whole cell samples and in a 60-70-nt form in OMVs (Fig. 2I). For both tRNA-Phe and CoA ligase northern blots, we observed a lower intensity signal and smaller sized RNA transcripts in OMVs as compared to whole cell samples indicating a differential loading of RNAs. Thus, our data suggest that ETBF and NTBF selectively load OMVs with a variety of transcripts that include full-length and partial transcripts.

### Outer Membrane Vesicles deliver RNA cargo to epithelial cell and co-localize with the endoplasmic reticulum

Having established strain-specific RNA signatures, we asked whether those RNAs reach the relevant cellular compartment in host epithelial cells. Using OMVs isolated from ETBF or NTBF, we first confirmed our co-staining of both RNA and OMVs using SytoRNASelect or Vybrant DiD, respectively (Fig. 3A; Supplemental Fig. 2). Using Caco2 cells incubated for 30 minutes with OMVs from ETBF, we demonstrate that OMVs with are taken up (Fig. 3B). To better understand where the OMV and RNA cargo were localizing, we stained the endoplasmic reticulum using a fluorescent antibody against protein disulfide isomerase (PDI) and treated the Caco2 cells with pre-stained OMV containing RNA, as described previously. Confocal microscopy images of these experiments demonstrate high co-localization (Mander’s split coefficient; average 0.85) of the OMV and RNA cargo with the ER (Fig. 3C/D). These results support the hypothesis that RNA cargo from *B. fragilis* OMVs is taken up by epithelial cells and delivered to the ER.

**Figure 3:**
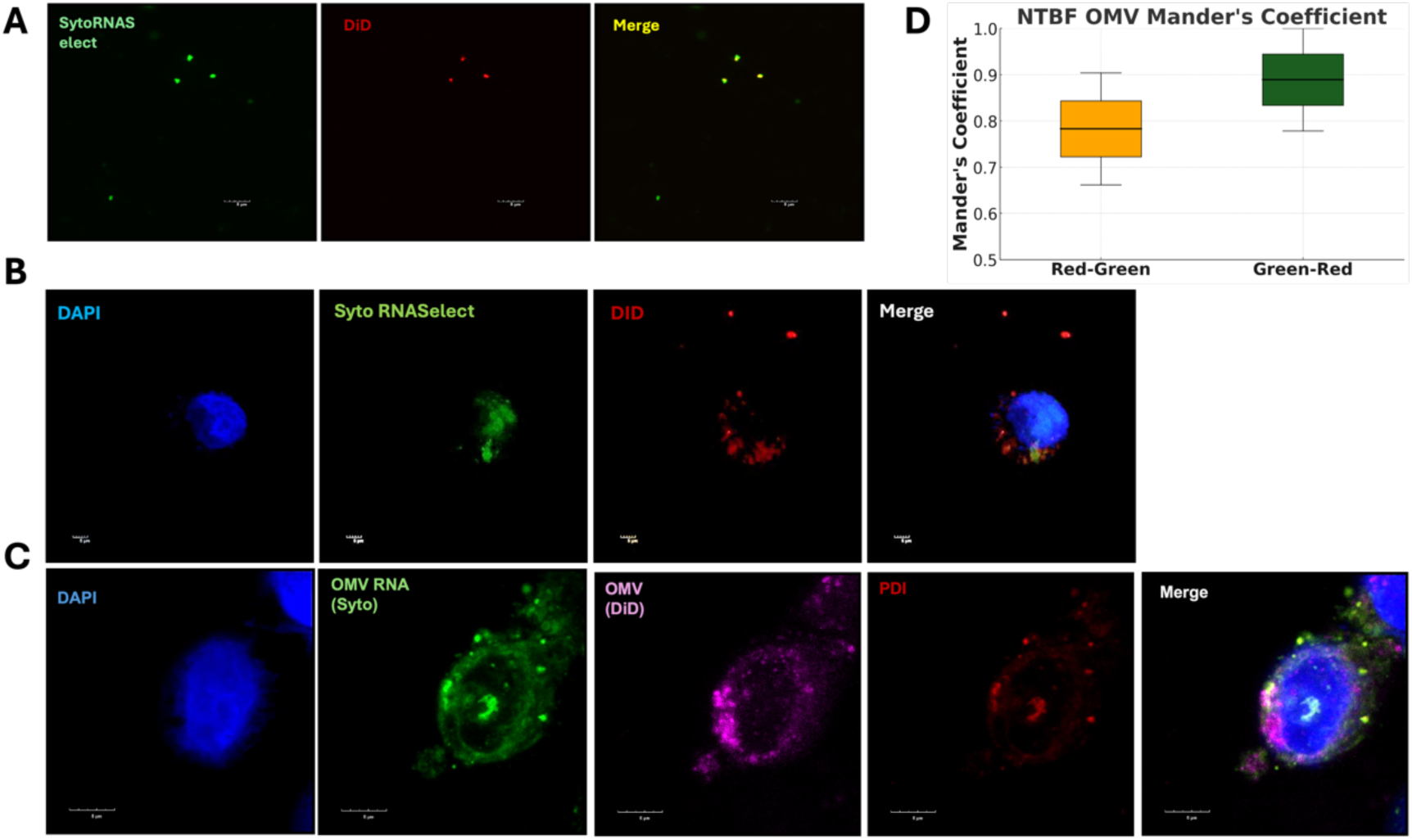
RNA cargo from ETBF and NTBF OMVs is taken up and localized to the endoplasmic reticulum in colonic epithelial cells. (A) Confocal microscopy images showing co-staining of RNA and OMVs using SytoRNASelect (green) and Vybrant DiD (red) in OMVs isolated from NTBF. The merged image indicates colocalization (yellow) of RNA and OMVs. Scale bar: 10 μm. (B) Confocal microscopy images of Caco2 cells incubated for 30 min with OMVs from ETBF. The images show the nuclei stained with DAPI (blue), RNA stained with SytoRNASelect (green), OMVs stained with Vybrant DiD (red), and the merged image showing colocalization (yellow) of RNA and OMVs within the cells. Scale bar: 10 μm. (C) Confocal microscopy images of Caco2 cells treated with pre-stained OMVs containing RNA, showing nuclei stained with DAPI (blue), OMV RNA stained with SytoRNASelect (green), OMVs stained with Vybrant DiD (magenta), endoplasmic reticulum (ER) stained with protein disulfide isomerase (PDI) (red), and the merged image showing colocalization of OMV RNA with the ER (yellow). Scale bar: 5 μm. (D) Boxplot depicting the normalized Mander’s split colocalization coefficients for images in (C) NTBF OMV and RNA cargo colocalization within colonic epithelial cells. The coefficients for Red-Green and Green-Red channels show significant colocalization in both channels.

### Specific OMV-associated RNAs are protected from RNase-treatment

To determine whether specific transcripts were contained within the OMVs and not simply co-purified, we treated intact OMVs with RNase prior to RNA extraction and quantified their abundances using qRT-PCR and northern blotting. First, we confirmed that our RNase treatment had a measurable impact on the total RNA concentration of our OMV samples (Fig. 4A), indicating that a portion of the RNA component of OMVs is susceptible to degradation. The ability to detect tRNA-Phe by northern blot was compromised by treatment with RNase (Fig. 4B), indicating that the tRNA-Phe could be present on the surface of the OMVs but might also be co-purifying during the OMV preparation from lysed cells. By contrast, the truncated RNA species of the CoA Ligase transcript remained detectable despite RNase treatment (Fig. 4C), indicating encapsulation within the OMVs or a specific modification that protects it from RNase degradation. We followed up on additional transcripts via qRT-PCR and note that the majority remained equally abundant, relative to rubredoxin, following RNase treatment (Sup. Table 1), suggesting encapsulation within OMVs.

**Figure 4:**
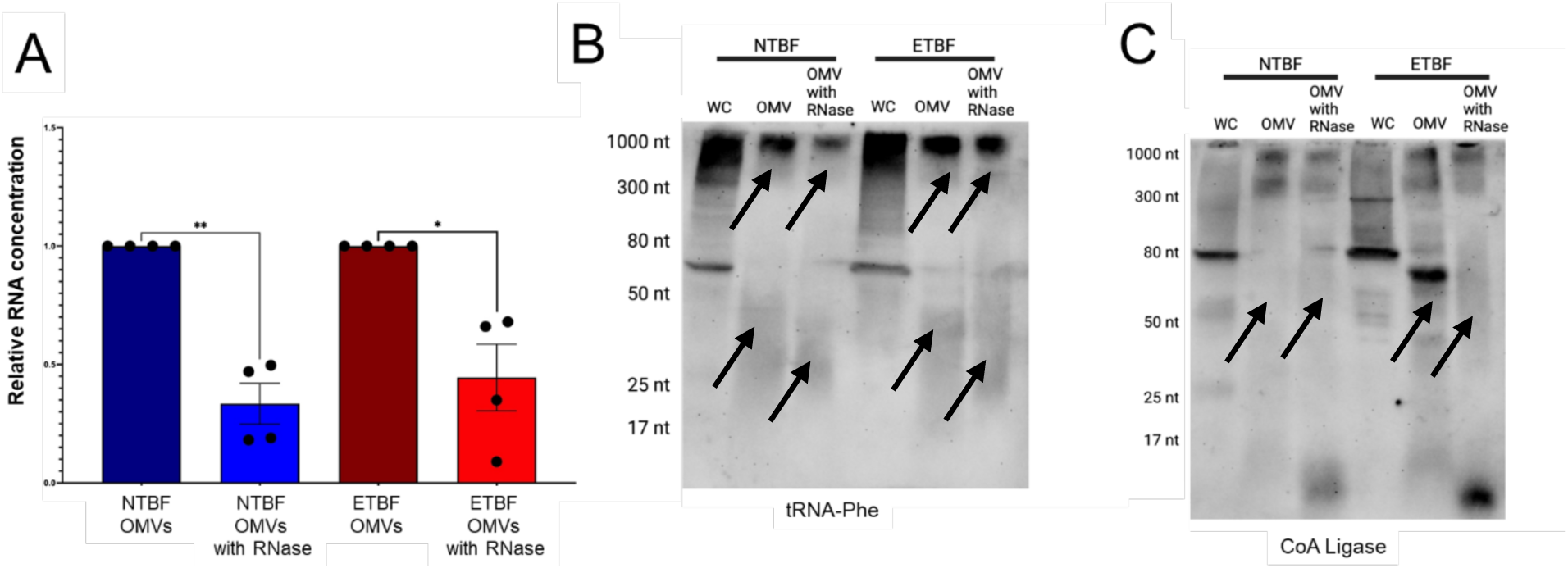
Specific OMV-associated RNAs are protected from RNase degradation. (A) Quantification of RNA recovered from equivalent amounts of OMVs with and without RNase treatment; individual points represent biological replicates (n=4) and data is presented as mean ± S.E.M.; statistical significance was determined using a Welch’s test. * p<0.05; ** p<0.01 (B/C). To confirm the size of tRNA-Phe (4B) and CoA Ligase (4C), northern blot probes complimentary to the most abundant RNAseq reads in the genes were used and successfully detected transcript fragments in RNase-treated OMVs (denoted by arrows). Probes bound to OMV-associated RNAs at shorter length in the in the blot compared to the whole cell samples, suggesting that the sequences contained in the OMVs are fragments of the larger RNA transcript.

### OMVs activate Toll-like Receptors in a dose-dependent manner

We next tested the functional consequence of the RNA profiles by measuring TLR activation in reporter cells. Innate immune receptors, including TLRs, respond to bacterial components, including macromolecules secreted within OMVs. We compared the capacity of ETBF and NTBF OMVs to activate TLR2 (which senses peptidoglycans), TLR3 (which recognizes dsRNA), TLR4 (which recognizes LPS), and TLR7 (which recognizes ssRNA). Treatment of reporter cells with either ETBF or NTBF OMVs showed strong activation of TLR2 at just 100 µg/ml (Fig. 5A), whereas activation of RNA-sensing TLR3 and TLR7 required 2000 µg/ml of OMV (Fig. 5B,C), and TLR4 activation was not observed (Fig. 5D). These data show a dose-dependent relationship on TLRs that. a primary response by non-RNA sensing at lower concentrations as opposed to RNA-sensing TLRs.

**Figure 5:**
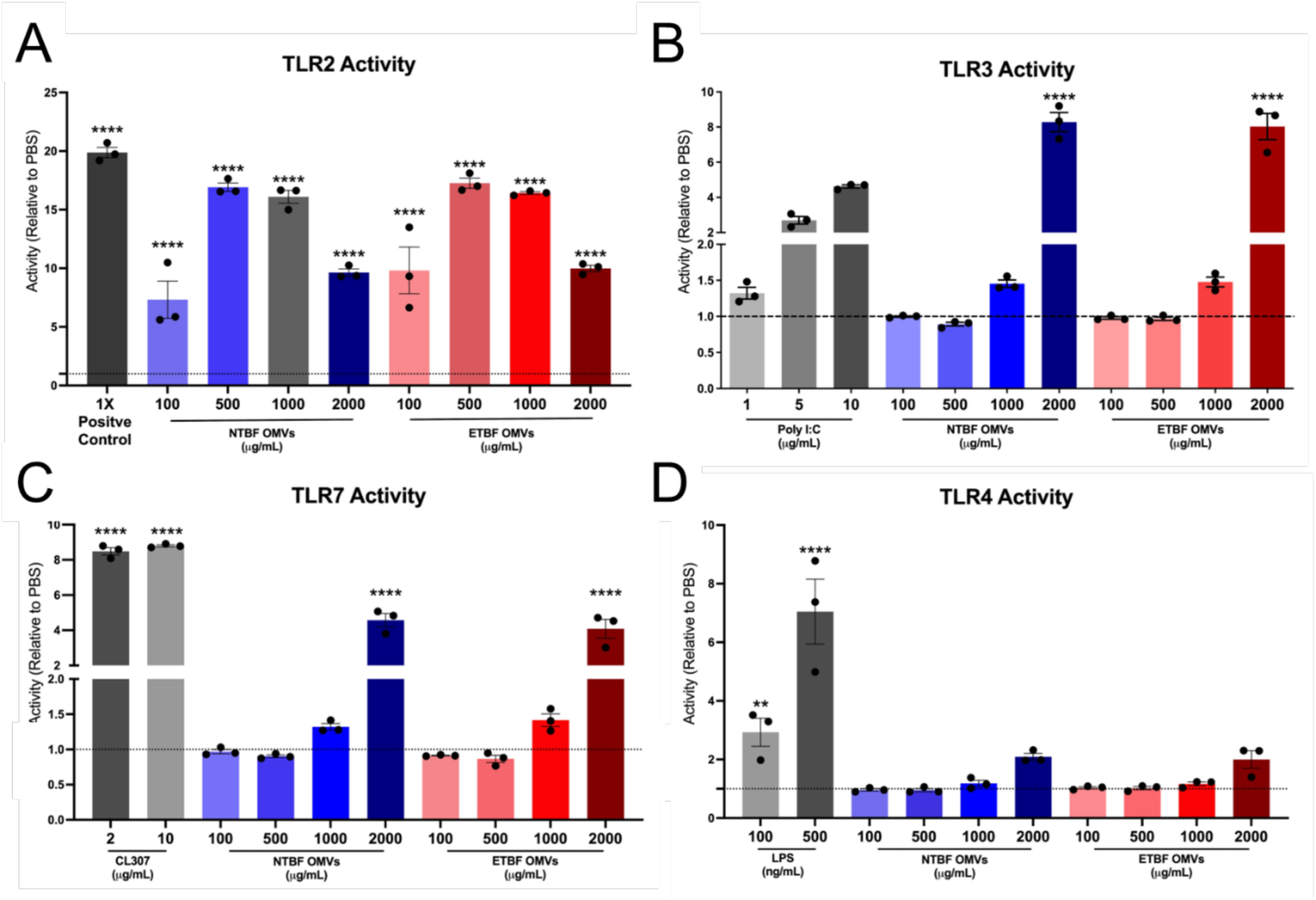
NTBF and ETBF OMVs activate TLRs in a dose-dependent manner. (A-D) OMVs from ETBF and NTBF were administered to HEK-293T cells expressing the indicated TLR proteins and an NFκB-responsive SEAP reporter gene. TLR activity was calculated by comparing vesicle and positive control treatments (gray columns) to the corresponding TLR cells treated with PBS (PBS treatment activity is represented by the dashed lines at y = 1). Points on the graph represent biological replicates (n=3) and data is presented as mean ± S.E.M. Statistical significance was found using Dunnett’s multiple comparison test. Stars represent significantly different from PBS control. ** p < 0.01, **** p < 0.0001. LPS = TLR4 agonist, Poly I:C = TLR3 agonist, CL307 = TLR7 agonist.

### OMVs from NTBF amplify the IL-8 response to TLR2 and TLR4 agonists in colorectal adenocarcinoma cells

The activation of various TLR receptors in the presence of NTBF and ETBF OMVs prompted us to investigate the role that the vesicles of different strains of *B. fragilis* have in contributing to immune signaling in colon cancer. To connect TLR activation to our key inflammatory read-out, we quantified IL-8 transcripts following OMV treatment, with or without OMV surface-RNA removal. For these experiments, we used two different colorectal adenocarcinoma cell lines: Caco-2 and HT29-MTX (Fig. 6). Caco-2 cells closely mimic the tight junction barrier of the colon, allowing the cell line to serve as a model to observe the interactions of the vesicles with host cells ^43^. HT29-MTX cells are a stable sub-population of the HT29 cell line that are differentiated into a goblet cell phenotype and can produce mucosal proteins that are found in the gastrointestinal tract ^44^. The distinct phenotypes of these cell lines represent distinct cell populations of the colon, allowing us to investigate the differential effects of vesicles on these cell types. We treated the cells for 2 hours with either 10 or 100 µg/mL of column-isolated vesicles, and we observed a significant induction of IL-8 expression in both cell lines in response to 100 µg/mL but not 10 μg/mL of both NTBF and ETBF OMVs (Supplemental Fig. 3A-B). Contrary to our initial hypothesis that OMVs from ETBF would elicit a greater response, marked by higher IL-8 expression, we observe a stronger response by high concentrations of OMVs from NTBF compared to ETBF. These results indicate a dose and strain-dependent response that distinguishes the commensal and pathogenic behavior of this species of bacteria on host inflammatory response.

**Figure 6:**
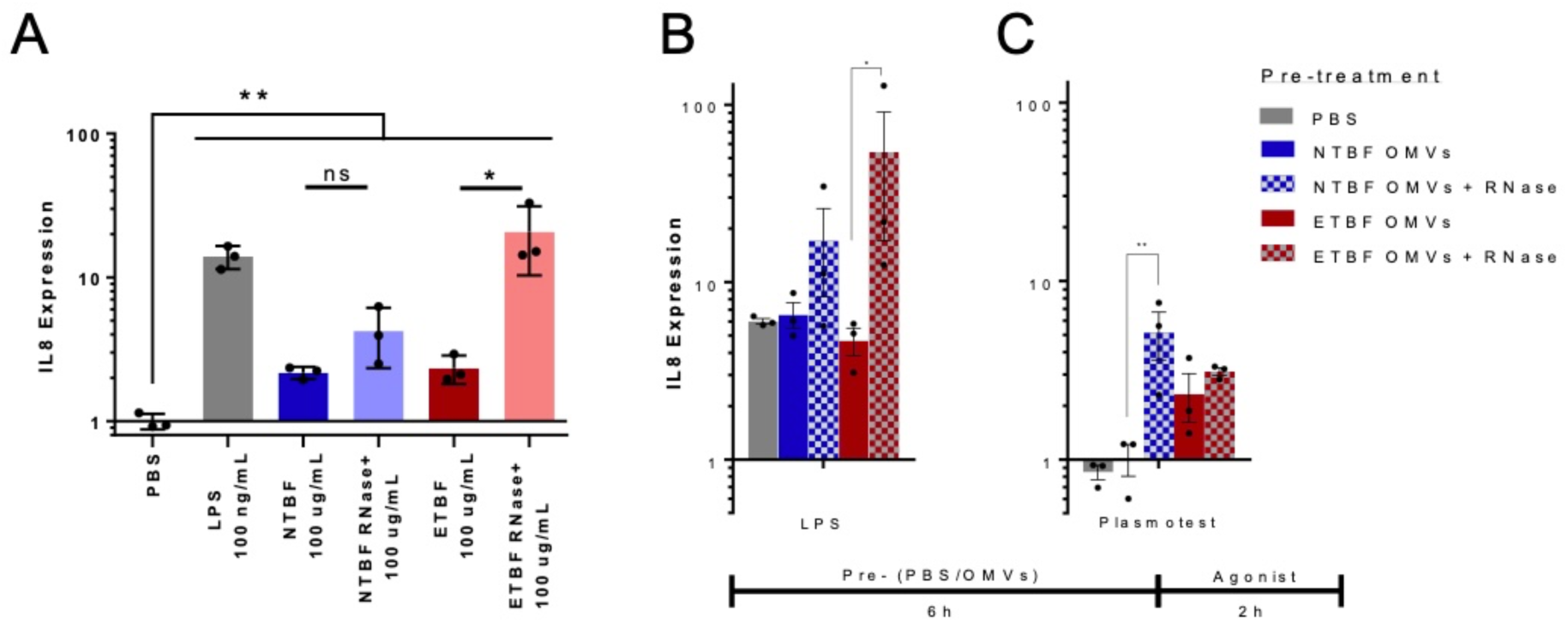
RNase-treated OMVs from ETBF amplify the IL-8 response to TLR2 and TLR4 agonists in colorectal adenocarcinoma cells. (A) Caco-2 cells treated with PBS, LPS, RNase-naïve or RNase-treated OMVs from NTBF and ETBF were assayed by qRT-PCR for IL8 expression. (B-C) Caco-2 cells treated with PBS, RNase-naïve or RNase-treated OMVs from NTBF and ETBF for 6h prior to withdrawal of pre-treatment and addition of LPS (TLR4 agonist) at 100 ng/mL (B) or Plasmotest (TLR2 agonist) (C). Signal normalized to vehicle only control. Points on the graph represent biological replicates and data is presented as mean ± st. dev. Statistical significance was found using Dunnett’s multiple comparison test. ** p < 0.01, *** p < 0.001, **** p < 0.0001.

### Extravesicular RNA suppresses IL-8 transcription and the capacity for TLR2 and TLR4 stimulation

Given that OMVs are capable of stimulating IL-8 expression, we next considered whether the RNase-accessible or RNase-protected RNAs mediated this effect. Thus, we tested whether removal of RNase-accessible RNAs from the OMVs would compromise the stimulatory effect of OMVs on IL-8 expression. Confirming the importance of RNase-protected RNAs, the capacity to stimulate IL-8 expression was not compromised in RNase-treated OMVs (Fig. 6C). Moreover, to our surprise, degradation of the RNase-accessible RNAs from ETBF OMVs yields vesicles with an *enhanced* stimulatory capacity (Fig. 6C). The same treatment of NTBF OMVs showed a trend towards enhanced stimulation though this was not statistically significant (Fig. 6C). To examine whether RNase-accessible RNAs affected the capacity of other TLR agonists to stimulate IL-8, we conducted a pre-treatment experiment. Loss of RNase-accessible RNAs led to a greater stimulatory capacity for TLR4 and TLR2 (significant for ETBF with a trend for NTBF). These results suggest to an immunosuppressive impact of RNase-accessible RNAs carried by ETBF and NTBF OMVs.

### Protein Components of B. fragilis OMVs

Finally, because OMVs also carry proteins that might modulate TLR responses and represent an additional biomarker for distinguishing differential effects between these strains, we compared the proteomic profiles of ETBF and NTBF OMVs. To characterize the differential loading of protein components by strains of *B. fragilis* into OMVs, we extracted total protein from OMVs and identified specific proteins using mass spectrometry. Analysis of electrophoretically separated proteins following silver staining shows distinctive bands between NTBF and ETBF OMVs (Fig. 7A), suggesting that OMV cargo differs between strains. We next sought to identify individual proteins that corresponded to high-intensity bands and which differed between ETBF and NTBF samples by extracting the proteins from gel fragments and analyzing by MS/MS. Assessing the confidently identified proteins that matched the molecular weight associated with the gel bands, five distinct proteins were identified in the ETBF samples and two in NTBF (Fig. 7B). Our results show that proteins identified in ETBF samples are primarily associated with carbohydrate metabolism, while proteins identified in NTBF samples are associated with hydrolase and DNA-binding activities (Fig. 7B). These distinct protein functions agree with previous studies ^45^, as NTBF and ETBF enzymes have been associated with carbohydrate metabolism; specifically, with hydrolases providing nutrition to NTBF and with energy-enriching metabolic pathways regulated by exported proteins in ETBF. ^24,46^ Thus, our results demonstrate similarities and differences in proteins identified between OMVs produced ETBF and NTBF.

**Figure 7:**
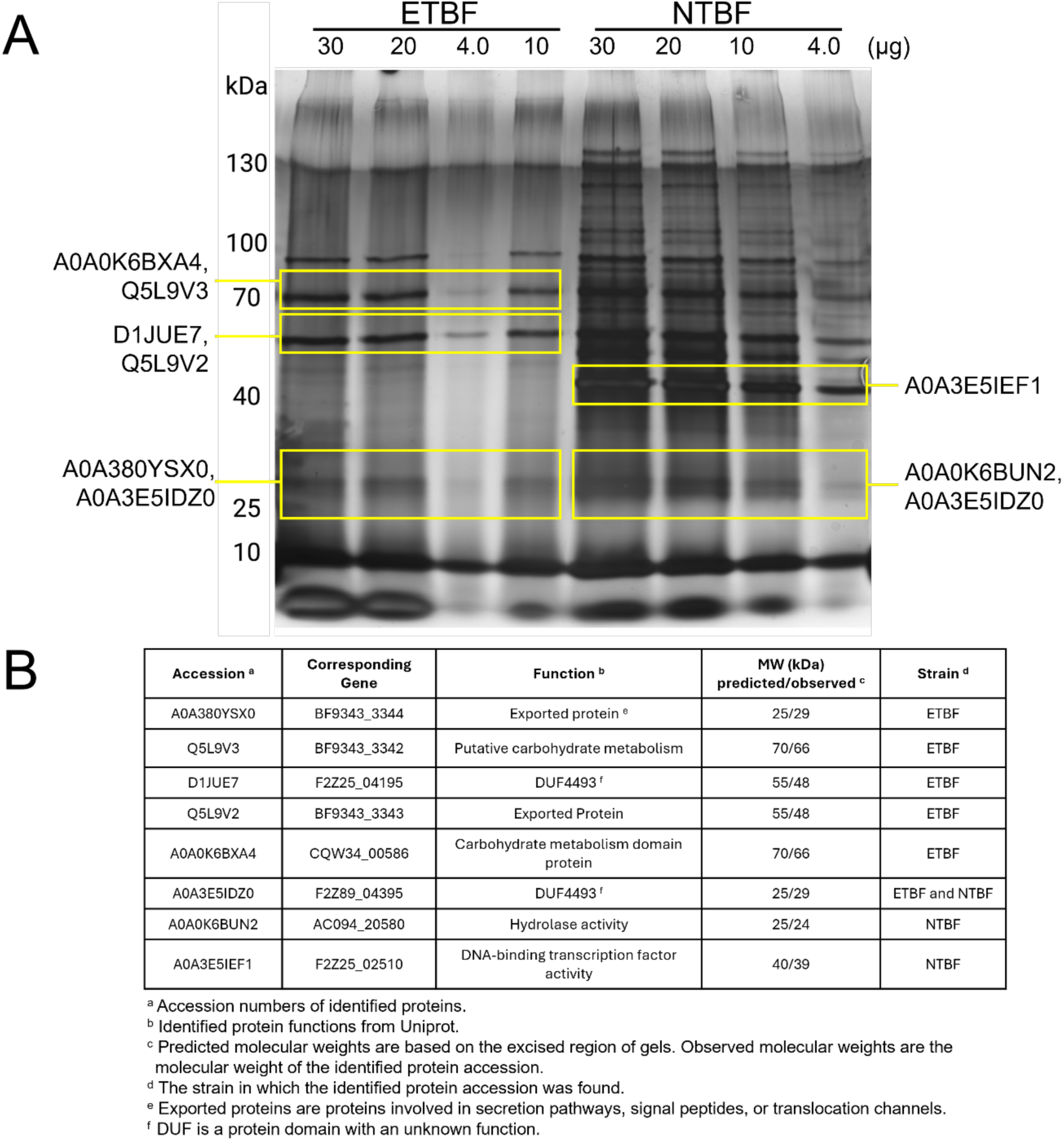
Protein components of *B. fragilis* OMVs differ between ETBF and NTBF. (A) Silver staining of OMV samples of ETBF and NTBF show different protein profiles. ETBF and NTBF OMVs at different concentrations were run on a 10% SDS-PAGE gel followed by silver staining. (B) Protein IDs detected for ETBF and NTBF with >1 peptide match and which possess a molecular weight within the range of excised gel band.

## Discussion

There is a growing interest in understanding the effects of bacteria-derived OMVs on host intestinal physiology and health ^23,29,47,48^. The immunogenic and immunosuppressive effects of OMVs on host cells and their ability to be easily altered to present specific antigens warrants further investigation of OMVs as a platform for therapeutic development and vaccine development ^49–51^. The effects of major biomolecular components of OMVs such as LPS, peptidoglycans, and secreted proteins (i.e. toxins) have been the subject of a majority of the research surrounding the characterization and downstream effects of OMVs on host cells ^23,34,52–54^. Comparatively, the role of nucleic acids (DNA and RNA) in OMVs, has only been recently begun to be characterized due to improvements in the isolation and high-throughput sequencing of OMV-derived nucleic acids^55^. The role of OMV-derived RNA in host physiology and disease has been described in common bacterial species that are clinically relevant such as *E. coli, P. aeruginosa, Vibrio cholerae,* and *P. gingivalis* ^25,39,56,57^. As researchers continue to delineate the role of the microbiome and of specific bacterial species on host health, it is imperative to understand the RNA profiles of their shed OMVs, since these clearly have important modulatory effects on downstream host pathways.

Towards this end, we sought to profile and characterize the RNA species contained within OMVs from two strain of *B. fragilis*, commensal (NTBF) and pathogen (ETBF), due to their distinct effects on chronic inflammation and colorectal cancer ^24,58–60^. Profiling the OMV-derived RNA reads to the general classes of gene coding regions of both genomes revealed an enrichment for protein coding-aligned RNA reads and a depletion of reads that aligned to tRNAs, rRNAs and pseudogenes. Many factors affect the distribution of RNA classes contained in OMVs, including stress, pH, phase of growth, and other culture conditions. In our experiments, we were careful to maintain consistency in all conditions throughout, including the optical density (OD) at which OMVs were isolated. In comparison to other studies, the protein-coding to non-protein-coding class can vary greatly. As demonstrated by Malabrirade *et al.*^61^, culture conditions can dramatically affect this ratio. In this study specifically, the majority of OMV RNA cargo was rRNA in control low OD, but decreased in the high OD samples. The size of the small RNA is also differentially abundant by RNA class, with large (>200 nt) RNAs being predominantly rRNA while smaller RNAs (<50 nt) making up a larger portion of tRNAs in *E. coli* and but not other bacteria ^56,62^. In studies of *A. pleuropneumonia*, OMVs tRNAs, and rRNA were most abundant, with tRNAs being enriched in OMVs compared to whole cells ^63^. Specifically, tRNAs were enriched in OMVs as compared to whole-cell RNA classes. Thus, overall, there are a variety of factors that affect the distribution of RNA types, which also vary by bacterial species; however, tRNA enrichment is a characteristic of OMVs from other bacterial species. The presence of these general categories of reads in proportions that are different in OMVs from their cells of origin suggests that bacteria have a molecular mechanism that directs more fragments of protein coding RNA reads into OMVs compared to other genic-aligned classes.

To further understand the differences in the RNA read profiles of OMV and whole cell samples, we performed differential expression analysis to identify specific genes and intergenic regions of interest that are enriched in each sample condition and in between strains. Using this dataset in conjunction with the read alignment count numbers, we identified several RNAs of interest for further validation. Other researchers have demonstrated the presence of specific OMV-associated RNA sequences using qPCR for genes that are known to be enriched in the vesicles of their species of interest ^39,56^. We were able to successfully detect transcripts of interest through qPCR and northern blotting using both OMV and whole cell samples. These data suggest that OMVs contain full-length and smaller fragments of the transcript, and that they are specifically loaded in the vesicles for extracellular transport.

To ensure that the detection of our target sequences was due to internalization of transcripts within vesicles and not from external presentation on the outer membrane or secretion into the culture broth, we treated OMVs with RNase. We found that there was a modest decrease in the concentration of RNA isolated from RNase-treated vesicles compared to their untreated counterpart, suggesting that a portion of the RNA we initially isolated were not contained within the vesicles, but also that some RNAs were loaded into the OMVs protecting them from RNase treatment. These findings correspond with other studies investigating RNA fragments in OMVs ^39^. *P. aeruginosa* has been reported to load a tRNA fragment into its OMVs for delivery to human epithelial cells where the fragment can act in a similar manner to miRNAs and down regulate IL-8 expression in the cells ^39^. The presence of RNA fragments in our NTBF and ETBF OMV samples may suggest a similar miRNA-like function of these sequences as the *P. aeruginosa* tRNA fragment and warrants further investigation.

To further understand the interactions of *B. fragilis* OMV-delivered RNA with host PRRs, we treated RNA-sensing TLR (TLR3 and TLR7) reporter cells with vesicles at various doses. TLR3 and TLR7 cells activated with treatments of OMVs at 2000 *μ*g/mL. The high concentration of OMVs required to activate these cells suggests that the RNAs contained in the vesicles may not be driving immune responses through TLRs. Nevertheless, vesicles of both strains induce expression of the pro-inflammatory cytokine IL-8. Other researchers have demonstrated that OMVs from commensal *E. coli* strains can induce *CXCL8* expression in Caco-2 and HT29 in a dose-dependent manner ^47^. These findings suggest a possible beneficial effect by priming the intestine to better respond to challenges from pathogen colonization as IL-8 is a chemokine that attracts neutrophils and macrophages to the affected site ^47,64^. Unexpectedly, when RNAs co-purified with OMVs are degraded by RNase, a significantly greater stimulation of IL8 transcription is achieved indicating a potential immunosuppressive role for these biomolecules. Notably, RNase-treated ETBF vesicles lose the 75 nt fragment of CoA ligase but retain the longer RNA species. RNA fragments originally transcribed from intergenic regions are also retained in RNase-treated vesicles. Whether specific gene products or classes of RNAs mediate these differential effects remains to be elucidated.

Altogether, our data demonstrate that NTBF and ETBF OMVs are enriched in specific RNA sequences from coding and intergenic regions of the *B. fragilis* genome. Through this work, we validated the presence of previously uncharacterized RNA sequences using qPCR and northern blotting, and we demonstrated that these sequences are contained within vesicles and differentially protected from RNase degradation. We further showed OMVs activate certain classes of TLRs, suggesting a role for these receptors in host-vesicle interactions and that ETBF and NTBF vesicles can produce strong inflammatory responses in epithelial colorectal cancer cells, dependent on the loading of intravesicular and extravesicular RNAs. Future research will be focused on understanding the biological implications of these small RNA cargo on host inflammatory responses, especially in the context of CRC. Overall, our research highlights a key factor, extravesicular RNAs, for understanding the role of OMVs on host immune responses through bacterial OMV RNAs.

## Acknowledgements

The authors wish to acknowledge the contributions of Dr. Cindy Sears for gifting *B. fragilis* strains, as well as Andrew Cox and Harry Ojeas for technical assistance.

---

**Supplemental Figure 1:**
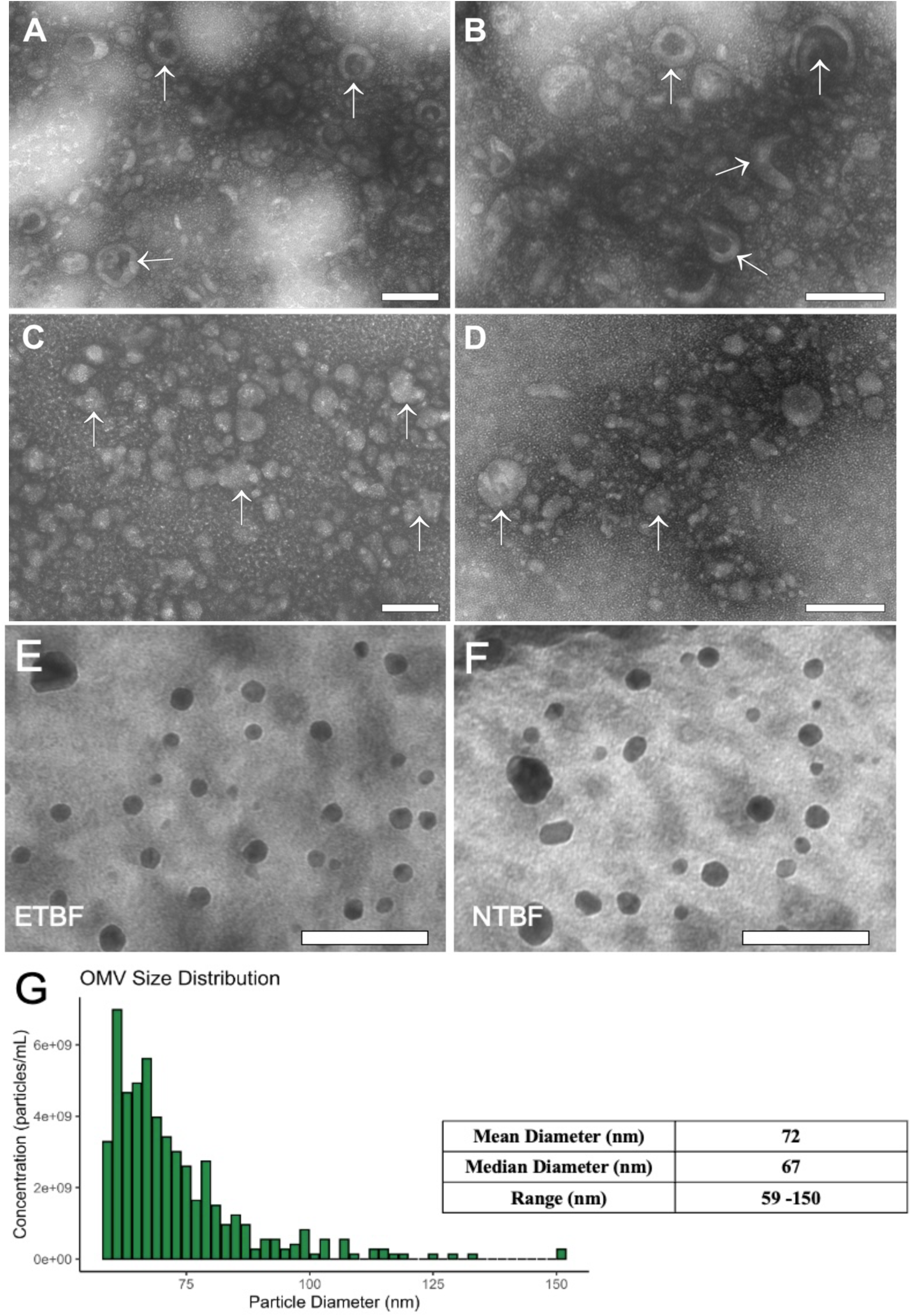
TEM imaging and size profiling of ETBF and NTBF OMVs. *Top Panel:* OMV staining with UA alone. TEM micrographs of ETBF (A and B) and NTBF OMVs (C and D) stained with uranyl acetate. Arrows denote vesicles that appear collapsed or which experienced a loss of structure. Scale bars, 100 nm. X 50,000 magnification. (Originally published in *Microscopy* as Fig. 1; Ref. 42) *Bottom Panel:* Concentrated samples of ETBF (E) and NTBF (F) vesicles were co-incubated with osmium tetroxide and uranyl acetate for imaging by TEM. (D) Quantification of ETBF OMVs in concentrated samples using nanoparticle tracking analysis (NTA). Scale bars, 100 nm. Magnification 50,000X

**Supplemental Figure 2:**
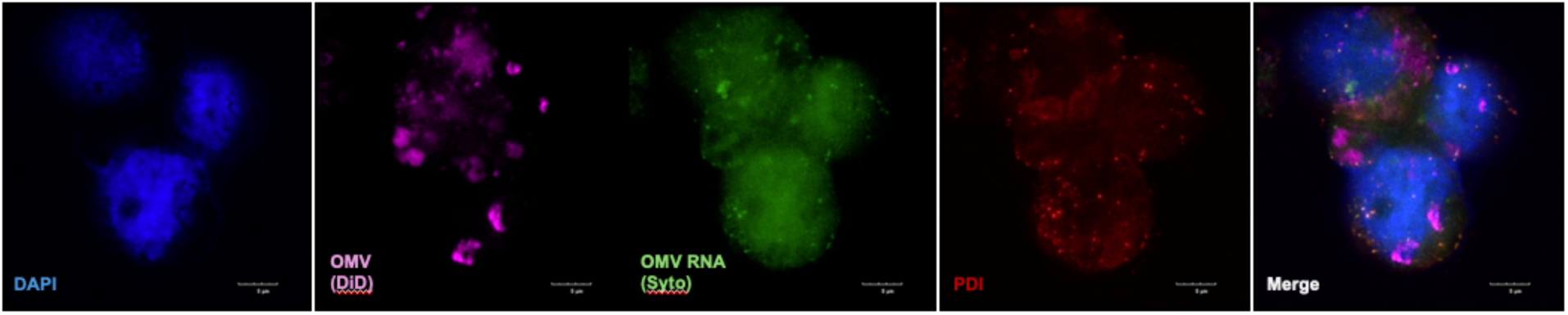
Additional confocal imaging showing OMV RNA localization to the endoplasmic reticulum in Caco2 cells. Confocal microscopy of Caco2 cells incubated for 1 hr with NTBF OMVs pre-stained with Syto RNASelect (green) and Vybrant DiD (purple). The endoplasmic reticulum (PDI) was labeled with Alexafluor 568 (red). Scale bar: 5 μm.

**Supplemental Figure 3:**
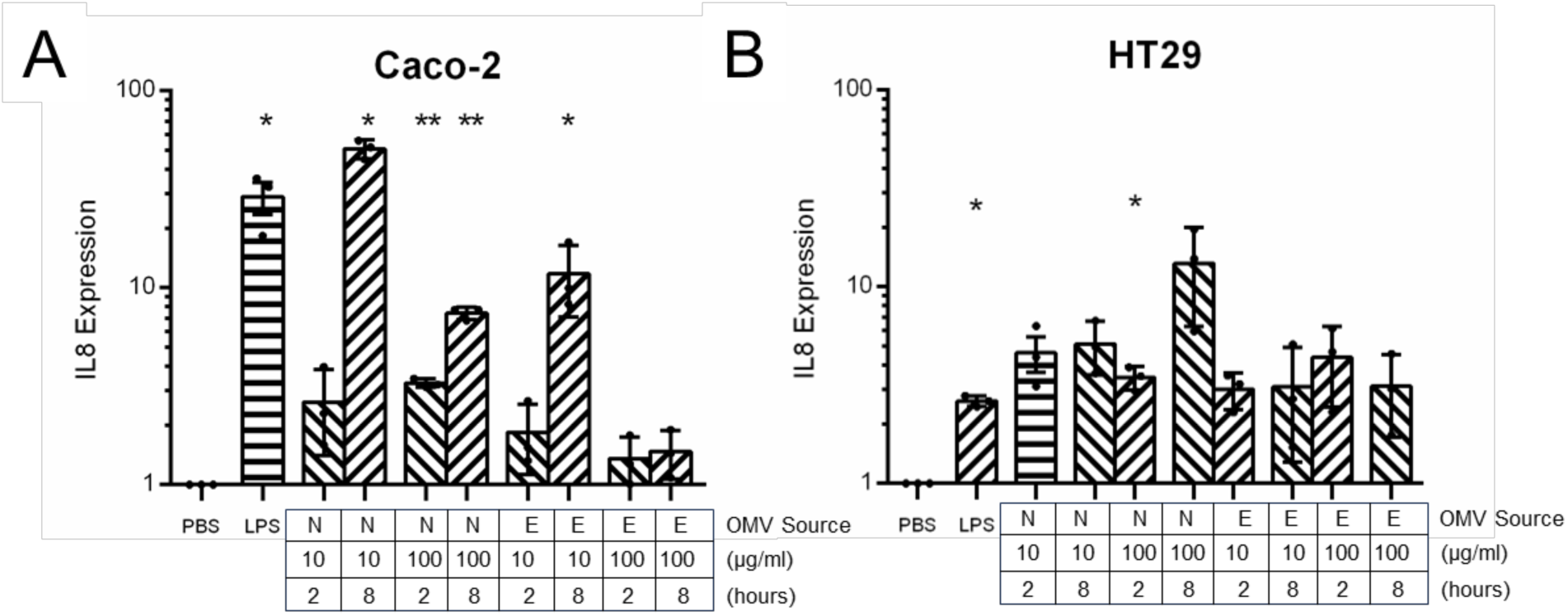
OMV dose-response effect on IL-8 expression in colon cancer cell lines. Caco-2 (A) and HT29-MTX (B) cells were treated with ETBF (E) and NTBF (N) vesicles for 2 or 8 hours prior to RNA extraction. qRT-PCR was performed for IL-8 transcript quantification. PBS and LPS were negative and positive controls, respectively. Dots represent biological replicates and data is presented as mean ± st. dev. Statistical significance was found using Dunnett’s multiple comparison test. ** p < 0.01, *** p < 0.001, **** p < 0.0001.

## Supplemental Materials and Methods

### Transmission Electron Microscopy

Concentrated OMVs were prepared for TEM imaging by staining with uranyl acetate and osmium tetroxide separately, then co-staining with osmium tetroxide and uranyl acetate. For uranyl acetate, copper grids were incubated on 10 *μ*L drops of OMVs for 5 minutes, followed by two washes on DI water drops for 2.5 minutes each and finally stained with 2% uranyl acetate for 1 minute. For the osmium tetroxide and uranyl acetate co-stain, 10 *μ*L of OMVs mixed with an equivalent volume of 4% osmium tetroxide for 10 minutes on copper grids followed by staining with 2% uranyl acetate for 1 minute. All grids had excess liquid blotted off with filter paper and were allowed to dry overnight. The grids were then imaged with a TEM (JEM-1010, JEOL Inc., Tokyo, Japan), a minimum of 7 images per test condition were obtained.

### Detailed Proteomics Methods

#### Materials

Dithiothreitol and LC-MS grade acetonitrile were purchased from Thermo Fisher Scientific (Waltham, MA). Proteomics grade iodoacetamide was from VWR International (Radnor,PA). Ammonium bicarbonate was purchased from Sigma Aldrich (St. Louis, MO). [Glu1]-Fibrinopeptide B (Glu-Fib) standard was purchased from Waters Corporation (Milford, MA). Nanopure water was obtained from a Purelab Flex 3 water purification system (Elga, Veolia Environment S. A., Paris, France).

#### Sample Preparation

Protein bands excised from an SDS gel with molecular weights of 25, 40, 55, 70, and 100 kDa were destained according to the manufacturer’s instructions (Pierce Silver Stain Kit, Thermo Scientific, 24612), and prepared individually for proteomic analysis. Proteins in gel pieces were reduced with dithiothreitol (1:10 protein-to-dithiothreitol molar ratio) on an Eppendorf ThermoMixer Temperature Control Device (Hamburg, Germany) for 30 minutes at 60 °C. Iodoacetamide (1:100 protein-to-iodoacetamide molar ratio) was added to alkylate free cysteines for 30 minutes at room temperature in the dark. Additional dithiothreitol (5:1 dithiothreitol-to-iodoacetamide molar ratio) was added to quench the alkylation reaction. Gel pieces were shrunk using 200 µL acetonitrile to cover the gel pieces and incubated for 15 minutes at room temperature. Acetonitrile was removed to waste, and gel pieces were air dried for 10 minutes. To digest proteins, trypsin/LysC (Promega, Madison, WI) was added (1:25 enzyme-to-protein ratio) with 100 mM ammonium bicarbonate (pH 8) and incubated on a ThermoMixer for 18 hours at 37 °C. The peptide-containing supernatant was moved to a clean sample vial for analysis. Remaining gel pieces were washed with 200 µL of 1 % formic acid in water for 5 minutes on a ThermoMixer to release any peptides trapped in gel pieces and the peptide-containing supernatant was added to the sample vial. The peptide mixture was dried down on an Eppendorf Vacufuge plus (Hamburg, Germany) for 30 minutes at 30 °C.

#### Liquid Chromatography – Tandem Mass Spectrometry

Peptides were diluted in 99 % water, 0.9 % acetonitrile, and 0.1 % formic acid; then injected onto a Waters nanoACQUITY UPLC with a Waters ACQUITY UPLC M-Class Symmetry C18 Trap Column (5 µm, 180 µm x 20 mm) coupled to a Waters ACQUITY UPLC PST BEH C18 nanoACQUITY analytical column (1.7 µm, 75 µm x 100 mm) (Milford, MA). A 36-minute gradient composted of solvent A (99.9 % water and 0.1 % formic acid) and solvent B (99.9 % acetonitrile and 0.1 % formic acid) was used at a flow rate of 0.5 µL/min: 0 min - 1 min, hold 1% B; 1 min - 30 min, increase B from 1% to 50%; 30 min - 31 min, increase B from 50% to 85%; 31 min - 36 min, hold 85% B; 36 min – 37 min, decrease B from 85% to 1%; 37 min - 45 min, hold 1% B.

Peptides eluting from the column were sprayed into a Waters Synapt G2-S (Milford, MA) using Fossilion Sharp Singularity LOTUS nESI emitters (Fossiliontech, Madrid, Spain). A data independent acquisition with a MS^E^ continuum method in positive-ion mode was used over a range of 250-1990 *m/z,* at a resolution of 20,000, and with a scan time of 0.6 seconds. Precursor ions were fragmented using collision-induced dissociation with a collision energy ramp from 20 V to 35 V. Glu-fib was used as the reference lock mass for post-acquisition mass correction and calibration.

#### Database Searching

Four sample-preparation replicates and three technical replicates were prepared for each excised band. Progenesis QI for Proteomics (version 4.2, Nonlinear Dynamics) was used to search replicates of each excised gel band against the reference proteomes: Uniprot Bacteroides Fragilis strain ATCC 25285 DSM 2151 CCUG 4856 JCM 11019 NCTC 9343, containing 5518 proteins, and Bacteroides Fragilis, containing 4645 proteins (downloaded July 2023). Imported data runs were corrected for Glu-1-Fibrinopeptide B lock mass and an ion accounting workflow was enabled. Search parameters consisted of peptide and fragment tolerances: automatic, enzyme specificity: cleavage of the C-terminus of lysine and arginine (/K-\P/R-\P), maximum ion charge: 20, missed cleavages: maximum 1, run alignment: automatic, peak picking sensitivity: automatic, maximum protein mass: 250 kDa. Modifications included fixed carbamidomethyl at cysteine, variable oxidation at methionine, and variable phosphorylation at serine, threonine, and tyrosine. Relative quantitation using Hi-3 with protein grouping was used to generate protein identifications. Confident protein identifications using a 1 % false discovery rate and ion matching requirements set to 5 fragments/protein, 2 fragments/peptide, and 1 peptide/protein were used. Protein identifications were filtered such that only identifications within ±10 kDa of the molecular weight of the respective excised gel bands were reported. Protein molecular weights were obtained from Progenesis QIP, and protein functions were extracted from Uniprot.

#### Data Availability

All LC-MS/MS raw data have been deposited into MassIVE and ProteomeXchange data repositories with the following accession numbers: MSV000095065 and PXD053204 for MassIVE and ProteomeXchange, respectively.

---

**Supplemental Table 1.**
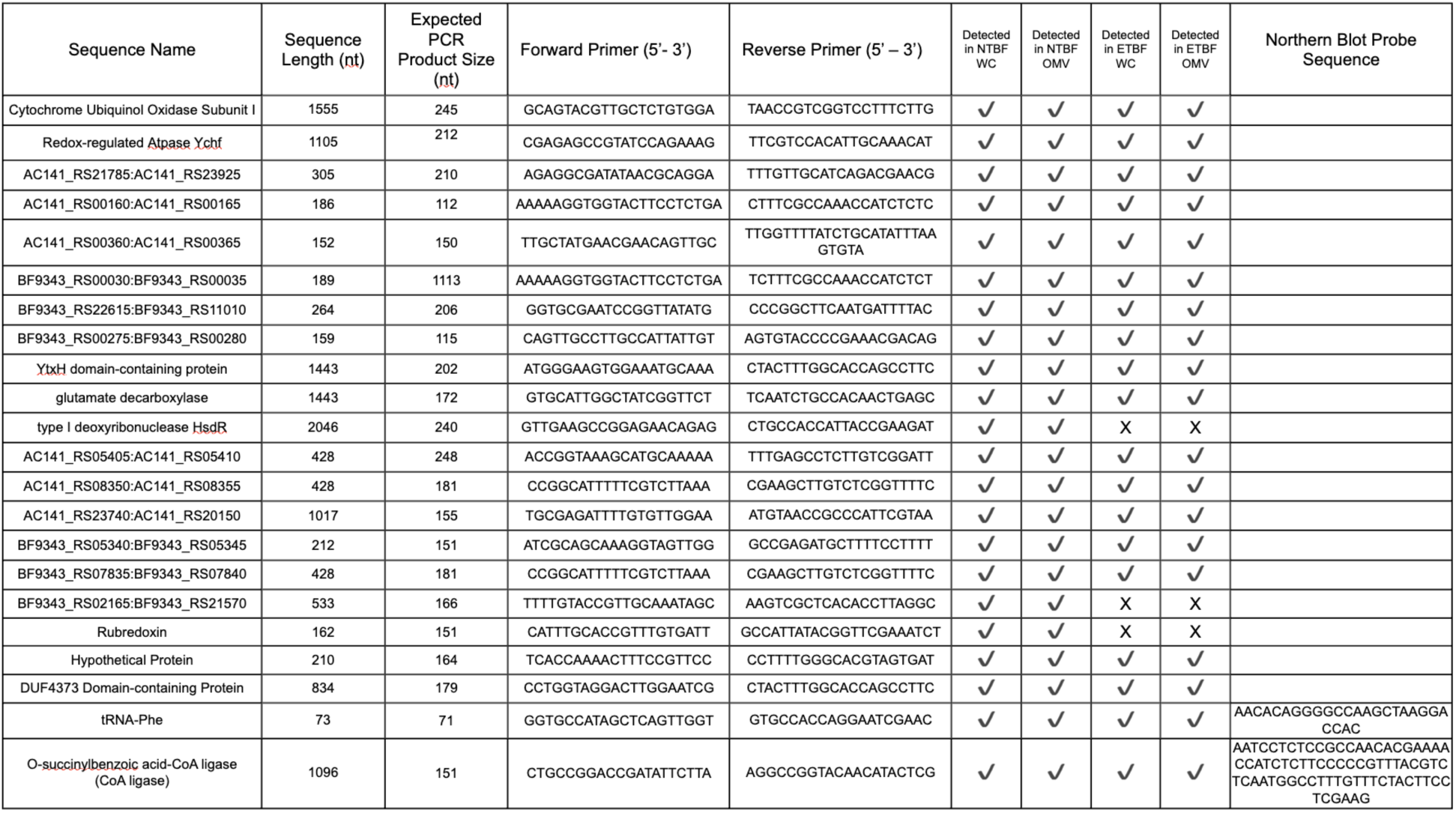
RNA species selected for analysis, along with PCR primers, northern blot probe sequences and a summary of qPCR results.

